# BioLitMine: advanced mining of biomedical and biological literature about human genes and genes from major model organisms

**DOI:** 10.1101/2020.07.17.208249

**Authors:** Yanhui Hu, Verena Chung, Aram Comjean, Jonathan Rodiger, Fnu Nipun, Norbert Perrimon, Stephanie E. Mohr

## Abstract

The accumulation of biological and biomedical literature outpaces the ability of most researchers and clinicians to stay abreast of their own immediate fields, let alone a broader range of topics. Although available search tools support identification of relevant literature, finding relevant and key publications is not always straightforward. For example, important publications might be missed in searches with an official gene name due to gene synonyms. Moreover, ambiguity of gene names can result in retrieval of a large number of irrelevant publications. To address these issues and help researchers and physicians quickly identify relevant publications, we developed BioLitMine, an advanced literature mining tool that takes advantage of the medical subject heading (MeSH) index and gene-to-publication annotations already available for PubMed literature. Using BioLitMine, a user can identify what MeSH terms are represented in the set of publications associated with a given gene of the interest, or start with a term and identify relevant publications. Users can also use the tool to find co-cited genes and a build a literature co-citation network. In addition, BioLitMine can help users build a gene list relevant to a MeSH terms, such as a list of genes relevant to “stem cells” or “breast neoplasms.” Users can also start with a gene or pathway of interest and identify authors associated with that gene or pathway, a feature that makes it easier to identify experts who might serve as collaborators or reviewers. Altogether, BioLitMine extends the value of PubMed-indexed literature and its existing expert curation by providing a robust and gene-centric approach to retrieval of relevant information.

## Introduction

Knowledge of biological systems exists in the form of millions of published literature citations in free text format. As of June 2020, there are about 31 million MEDLINE records, dating back to 1781, and accumulation of new publications has increased exponentially in recent years. The National Center for Biotechnology Information (NCBI) PubMed search engine for the MEDLINE database has been a fundamental resource, allowing scientists and physicians to identify publications of interest in the biological and biomedical literature. Nevertheless, many literature mining tools, often called “PubMed derivatives,” have been developed to facilitate the search of literature to address specific needs.

PubMatrix, for example, uses text mining to build frequency matrix of term co-occurrence (Becker *et al*. 2003). Similarly, MedGene builds gene and disease relationships based on co-occurrence of medical subject heading (MeSH) disease terms and genes (Hu *et al*. 2003). SciMiner is a web-based literature mining and functional analysis tool that identifies genes and proteins using a context-specific analysis of MEDLINE abstracts and full texts based on the list of PubMed publications provided by the user (Hur *et al*. 2009). EBIMed (Rebholz-Schuhmann *et al*. 2007) and GLAD4U (Jourquin *et al*. 2012) are gene-centric text mining tools analyzing the features for proteins/genes from MEDLINE and help to build gene list relevant to diseases as well as key words of biological concepts. PubTator (Wei *et al*. 2013; Wei *et al*. 2019) and Textpresso (Muller *et al*. 2004; Muller *et al*. 2018) are tools for viewing and retrieving bio-concept annotations in full text biomedical articles, facilitating literature curation using an automating text mining process.

Unlike PubTator and Textpresso, which were developed to mine the subset of PubMed records with full literature available, BioLitMine was developed to mine all MEDLINE records and to expand the features of some of the existing tools by providing a user-friendly web interface and visualization of search results, such as a word cloud of gene-to-MeSH search results and a network view of co-cited gene search results. We were also inspired by the idea behind GeneMatcher (Sobreira *et al*. 2015), a system that connects human geneticist to the scientists working on the corresponding orthologous genes in model organisms. However, different from the submission-based approach used by GeneMatcher, for BioLitMine we processed author information from PubMed literature and build gene-to-people relationships from all literature relevant to a gene, such that users can easily find all principle investigators associated in the literature with a given gene of interest. In addition, we also imported annotations of major signaling pathways to build pathway-to-people relationships from the literature so that users can easily find relevant investigators, genes, and publications. Altogether, BioLitMine serves as a useful platform for gene-centric mining of the published literature and other applications.

## Results and Discussion

### The BioLitMine pipeline and database

We recognized that existing associations of the PubMed literature with genes and medical subject heading (MeSH) terms, which include anatomy, disease, and other types of terms, could serve as the basis for a robust new literature-mining resource. To accomplish this, we built the BioLitMine pipeline, database, and user interface using a typical three-layer structure with a front end web-based user interface and back-end MySQL database that stores the literature information, as well as pre-computed analyses such as gene-to-mesh and gene-to-people relationships (**Fig. 1** and see ***Methods***). We rely on NCBI Pubmed “gene2PubMed” release files for association of genes with publications. Moreover, the association of publications with genes allows us to further associate publications (and their authors) with pathways, as well as to map from one species to another based on ortholog relationships.

**Figure 1:**
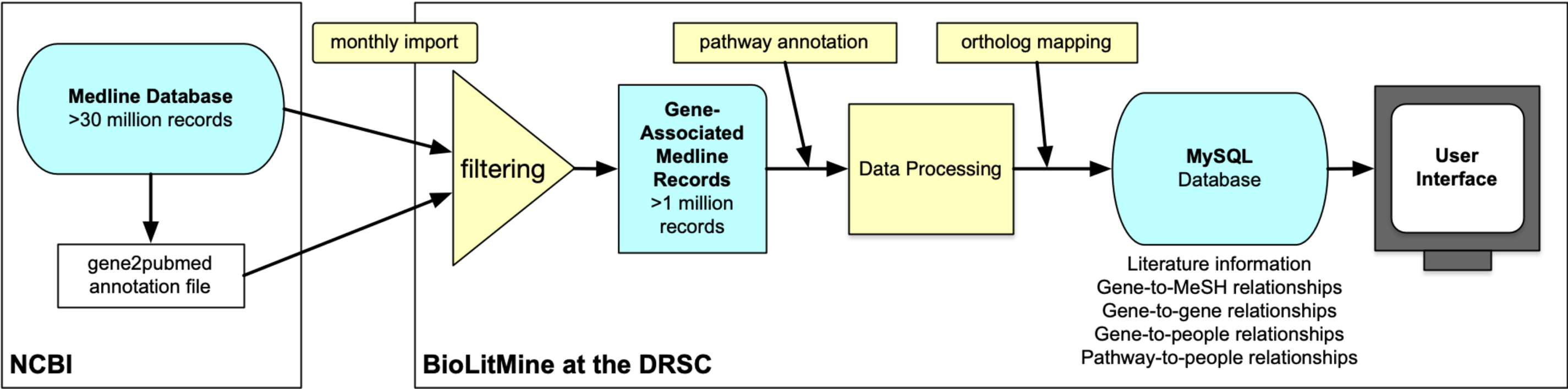
The BioLitMine literature mining and annotation informatics pipeline. Literature and gene associations are imported monthly and automatically from the MEDLINE database and NCBI PubMed release files. Literature records are then filtered to remove publications not associated with any gene. The resulting set of gene-associated records are then associated with pathway annotations and processed. DIOPT ortholog mapping is used to associate genes with orthologs in other species. All information is imported into a MySQL database and presented to users via an online user interface that facilitates search and retrieval using gene, MeSH term, people, and other information.

The pipeline we built first processes citations from the NCBI MEDLINE database, then, at the beginning of each month, automatically updates information stored in a MySQL database, minimizing the need for maintenance by developers while at the same time ensuring an up-to-date resource. New publications and gene-to-paper assignments are updated daily by PubMed. Our pipeline retrieves new MEDLINE incremental release files and gene2pubmed annotation files at each update. The pipeline then selects gene-relevant papers from all release files, including baseline files, and processes the associated information. Pathway annotations are retrieved from GLAD (Hu *et al*. 2015), SignaLink (Csabai *et al*. 2018), and KEGG (Kanehisa and Goto 2000). Ortholog mapping is done using the DRSC Integrative Ortholog Search Tool (DIOPT) ortholog mapping resource (Hu *et al*. 2011).

As of June 2020, our BioLitMine pipeline has processed 1,332 MEDLINE release files of 327 GB, including 1,015 baseline files released in December 2019 and 217 daily update files. Of the 31 million MEDLINE records available as of summer 2020, about 1.26 million records are annotated as related to genes. Of these, 1.13 million records are related to genes from the nine major model organisms and human included at BioLitMine. We note that gene-relevant publications account for 2% of publications in the period 1980-1999 and 7% of publications in the most recent 20 years (2000-2019).

### BioLitMine supports robust gene-based searches of the published literature

BioLitMine was designed to support gene-based search of PubMed records in a way that obviates problems associated with synonymous gene names by relying not on the genes as reported in the papers but instead, on gene2 pubmed associations annotated by NCBI. For a given gene input, BioLitMine first retrieves MeSH terms associated with the gene in the literature and displays the results in a table format (**Fig. 2A**). The user can then opt to view the search results as a ‘word cloud’ in which the size of each MeSH term is scaled based on the number of publications. Users can quickly identify a subject(s) of the interest and retrieve corresponding PubMed records. Alternatively, users can start with a given input gene of interest and identify co-cited genes and associated papers (**Fig. 2B**). After display of tabular results of the search, users can view a co-citation network that is built and compared with protein-protein interactions, genetic interactions and interolog annotations from the molecular interaction search tool (MIST) (Hu *et al*. 2018) (**Fig. 2B**, overlay panel). Finally, the “Batch Search” option at BioLitMine supports identification of publications associated with each gene on a list (**Fig. 2C**).

**Figure 2:**
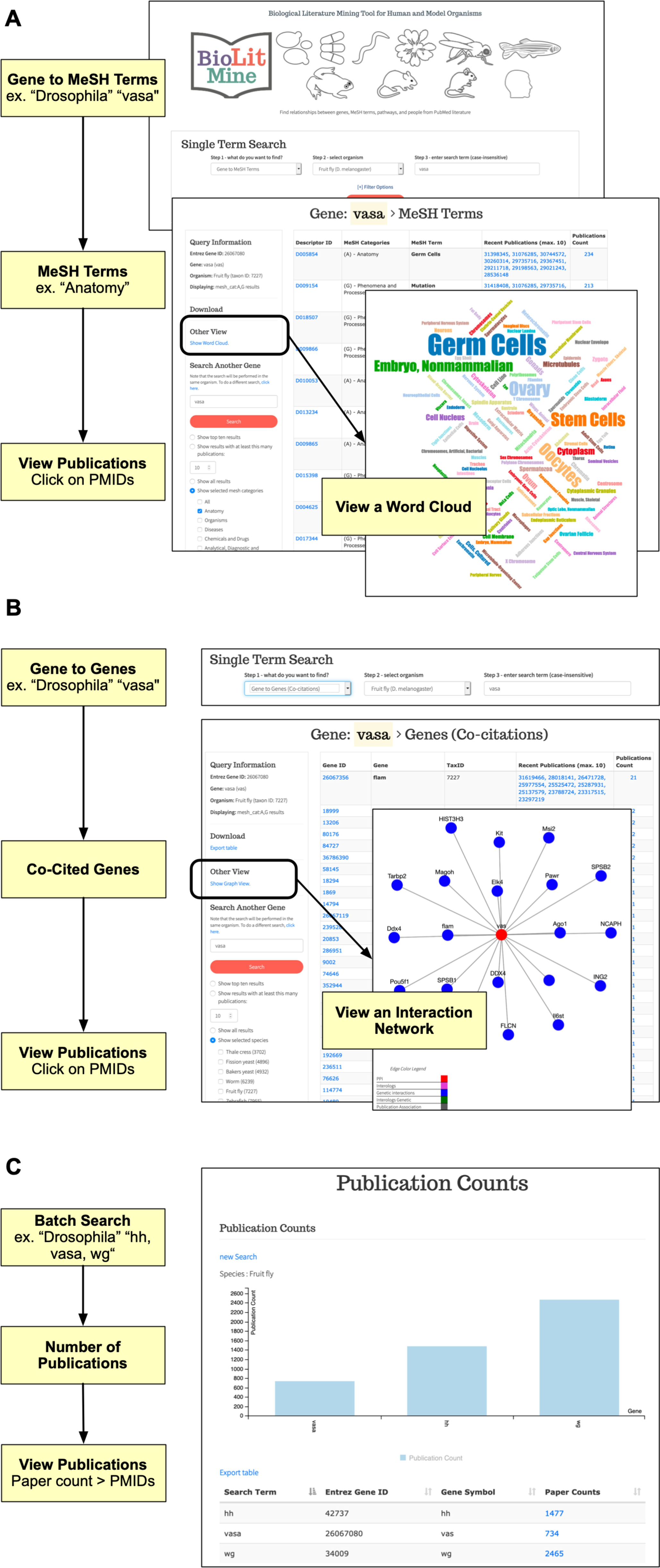
BioLitMine gene search. **A**. Example gene-to-MeSH term workflow using “Drosophila” as the selected species and “*vasa*” as the input gene (top panel). MeSH terms associated with the input gene are displayed and can be filtered to select a specific sub-category of terms (middle panel). In this example, only “anatomy” MeSH terms are displayed. Associated publications are provided as active links to records at NCBI PubMed. Users can also click to view the results in an alternative ‘word cloud’ visualization (overlay panel) and can download tabular results as a comma-separated value (csv) file. **B**. Example search for genes co-cited with an input term. As for (A), “Drosophila” was selected as the species and “*vasa*” as the input gene (top panel). Tabular results display genes co-cited with the input gene (bottom panel). Associated publications are provided as active links to records at NCBI PubMed. Users can also click to view the results in an alternative network visualization format (overlay panel) and can download tabular results as a csv file. **C**. Multiple genes can be searched using the Batch Search option. As shown in the example output, a count of publications is shown for each gene and links to the associated PubMed records.

### BioLitMine supports MeSH term-based search of genes and literature

For any MeSH term, BioLitMine first retrieves all relevant papers and then uses the pubmed2gene associations to extract genes associated with that subset of publications. The results are displayed as a table and are sorted based on the count of relevant publications, with the term associated with the largest number of publications displayed first (**Fig. 3**). Because MeSH terms are organized in hierarchical structure, users have the option to include more specific ‘child’ terms associated with the input term. For example, a search initiated using the term “Breast Neoplasm” will retrieve child terms, such as “Breast Carcinoma in Situ” and “Triple Negative Breast Neoplasms,” which can either be included or excluded in the gene search. After terms are defined, genes associated with the search are displayed (**Fig. 3**). In addition to retrieving genes for the organism selected in the initial search, BioLitMine also gives users the option to view a corresponding gene list for a different model organism (**Fig. 3**, bottom overlay panel) using DIOPT-based ortholog mapping (Hu *et al*. 2011).

**Figure 3:**
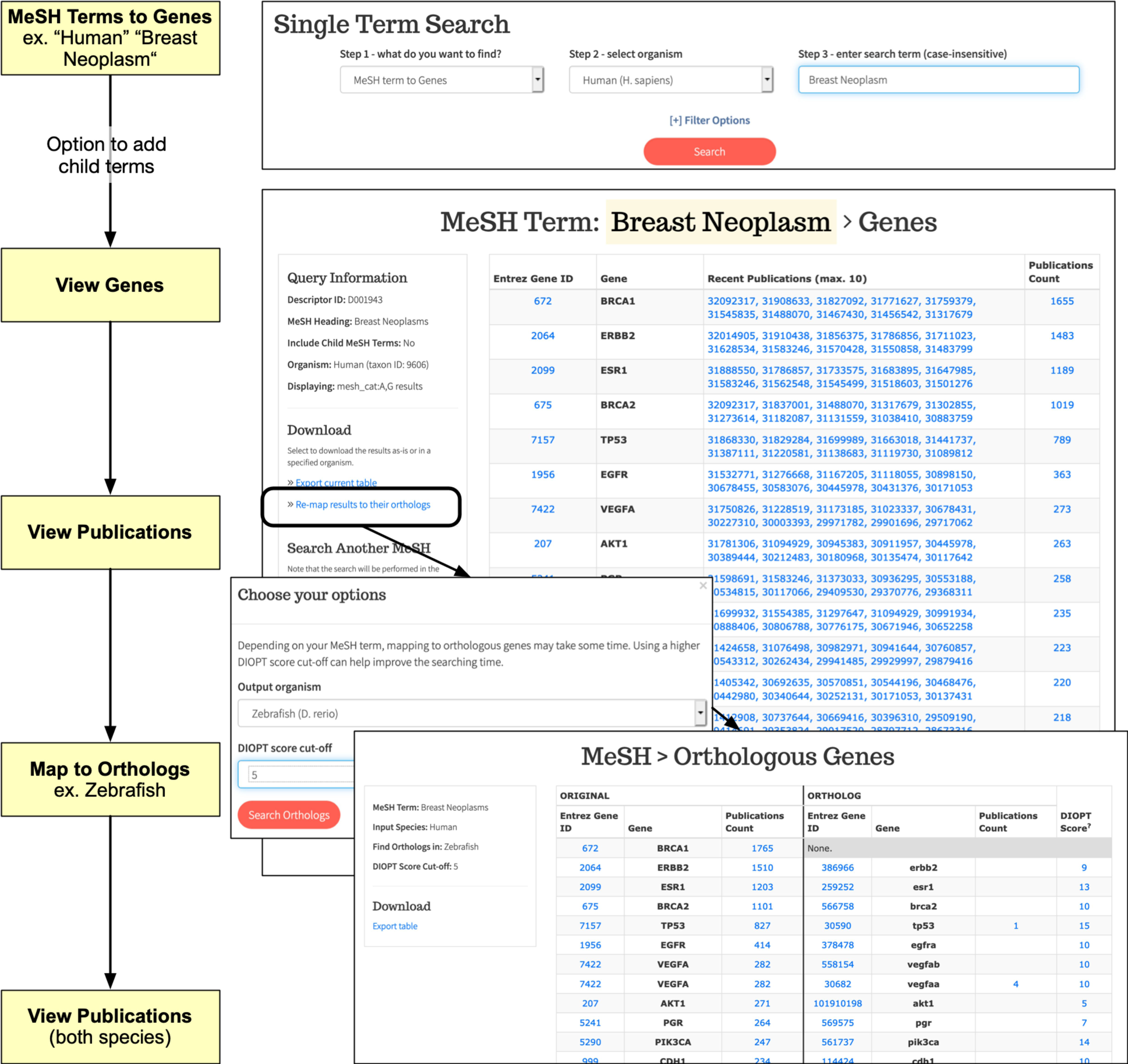
BioLitMine medical subject heading (MeSH) search. Example MeSH term-to-gene search using “human” as the selected species and “breast neoplasm” as the input term (top panel). Users are first given the option to choose ‘child’ terms additional to the input term (not shown). Next, a table of results is displayed. Associated publications are provided as active links to records at NCBI PubMed. Users can download tabular results as a csv file. In addition, the resulting gene list can be mapped to orthologs in another species (bottom overlay panels; “zebrafish” in this example).

### BioLitMine supports identification of authors associated with genes or pathways

We were interested to explore the possibility of using the published literature to identify experts, including experts based in a given region, in an automated manner. To support this, BioLitMine captures the last authors of all publications relevant to each gene and makes it easy for users to find experts for a given gene. The search results display the name of the last author, the number of publications associated with that author, and the year and address from the most recent publication. The results table can be sorted differently, e.g. by the number of papers or by the year of the most recent publication. In addition, we also imported annotation of major signaling pathways from GLAD (Hu *et al*. 2015), SignaLink (Csabai *et al*. 2018), and KEGG (Kanehisa and Goto 2000) so that users have the option to search people based on pathway annotations (**Fig. 4**). In addition to all of the fields displayed for gene-to-people search results, for a pathway2people search, the number of genes studied is also displayed and sortable. There is a possibility, although small, that some scientists who have the same last and first names might study the same pathway or even the same gene. This could be recognized by users by looking at the relevant literature, which is linked to in results tables. The problem might be obviated in the future by adoption at PubMed of unique identifiers for authors, such as ORCID IDs (Bohannon and Doran 2017).

**Figure 4:**
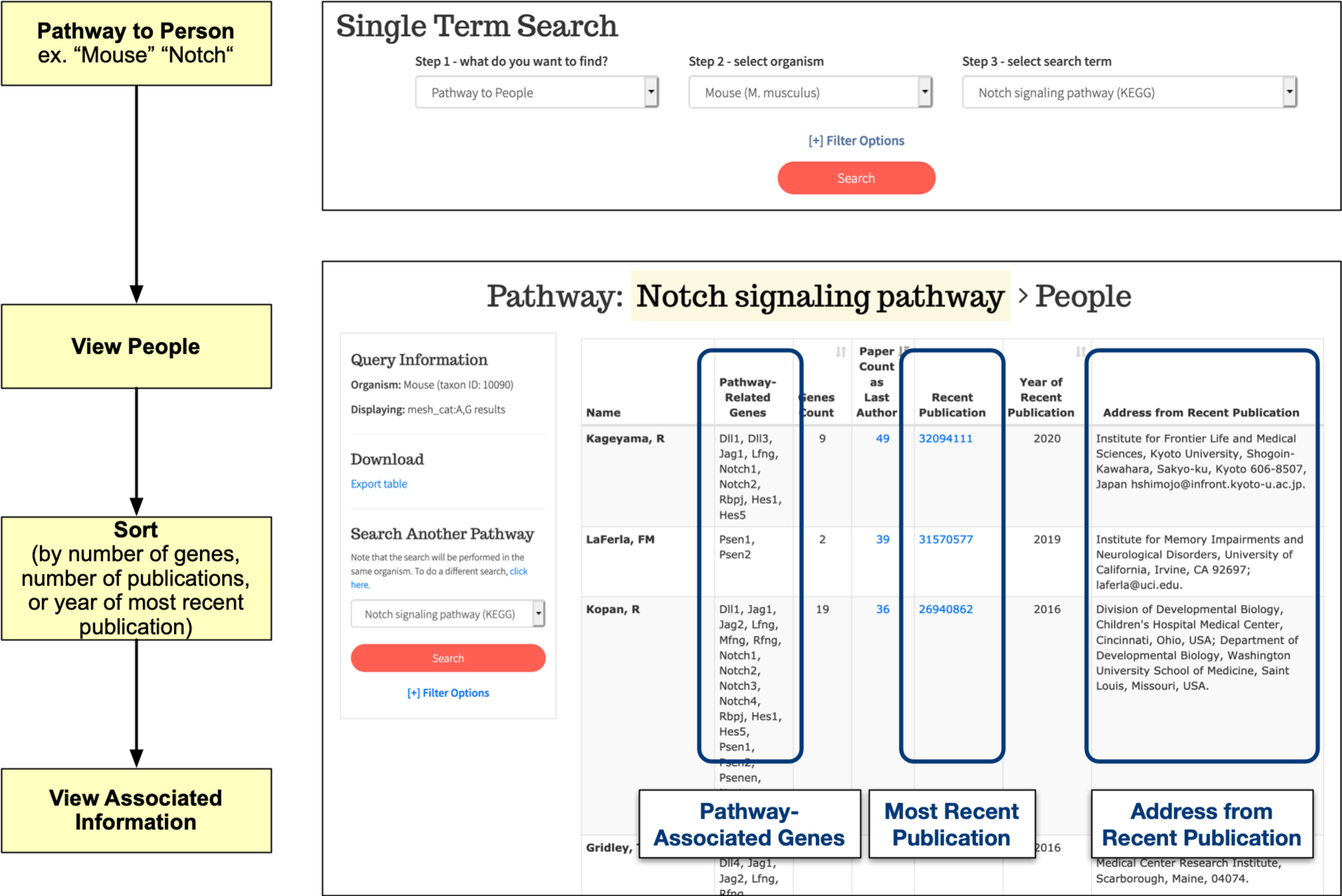
BioLitMine people search. Example of a pathway-to-people search using “mouse” as the selected species and “Notch” as the selected pathway (top panel). Gene-to-people searches are also supported (not shown). Results are shown in a table (bottom panel) that includes a list of genes associated with both the pathway and the person (last author); the count of genes and publications associated with both the pathway and person; a link to the most recent relevant publication and the year of that publication; and an address extracted from the most recent publication. Results can be downloaded as a csv file.

### BioLitMine supports MeSH term enrichment analysis

In addition to batch searches with a gene list to retrieve publications (**Fig. 2C**), BioLitMine also provides an option for users to input a gene list and then perform an enrichment analysis to identify MeSH terms over-represented in the set of related publications (**Fig. 5**). To test the enrichment feature, we analyzed MeSH terms in the “Phenomena and Processes” category after uploading the set of 208 *Drosophila* genes annotated as autophagy-related genes at GLAD (Hu *et al*. 2015) and compared the top 25 MeSH terms retrieved at BioLitMine with the top 25 GO terms retrieved using DAVID (https://david.ncifcrf.gov/) (Jiao *et al*. 2012). As shown in **Table 1**, we found that the majority of biological process terms are consistent between MeSH term enrichment by BioLitMine and GO-term enrichment by DAVID. For example, GO enrichment results identify “cellular responses” whereas MeSH term enrichment identifies “glycogenolysis/lipid metabolism” as relevant to the gene list. GO enrichment analysis is widely used for analyzing gene lists and BioLitMine provides a complimentary approach, covering biological themes that are not annotated or not fully annotated by gene ontology.

**Table 1.**
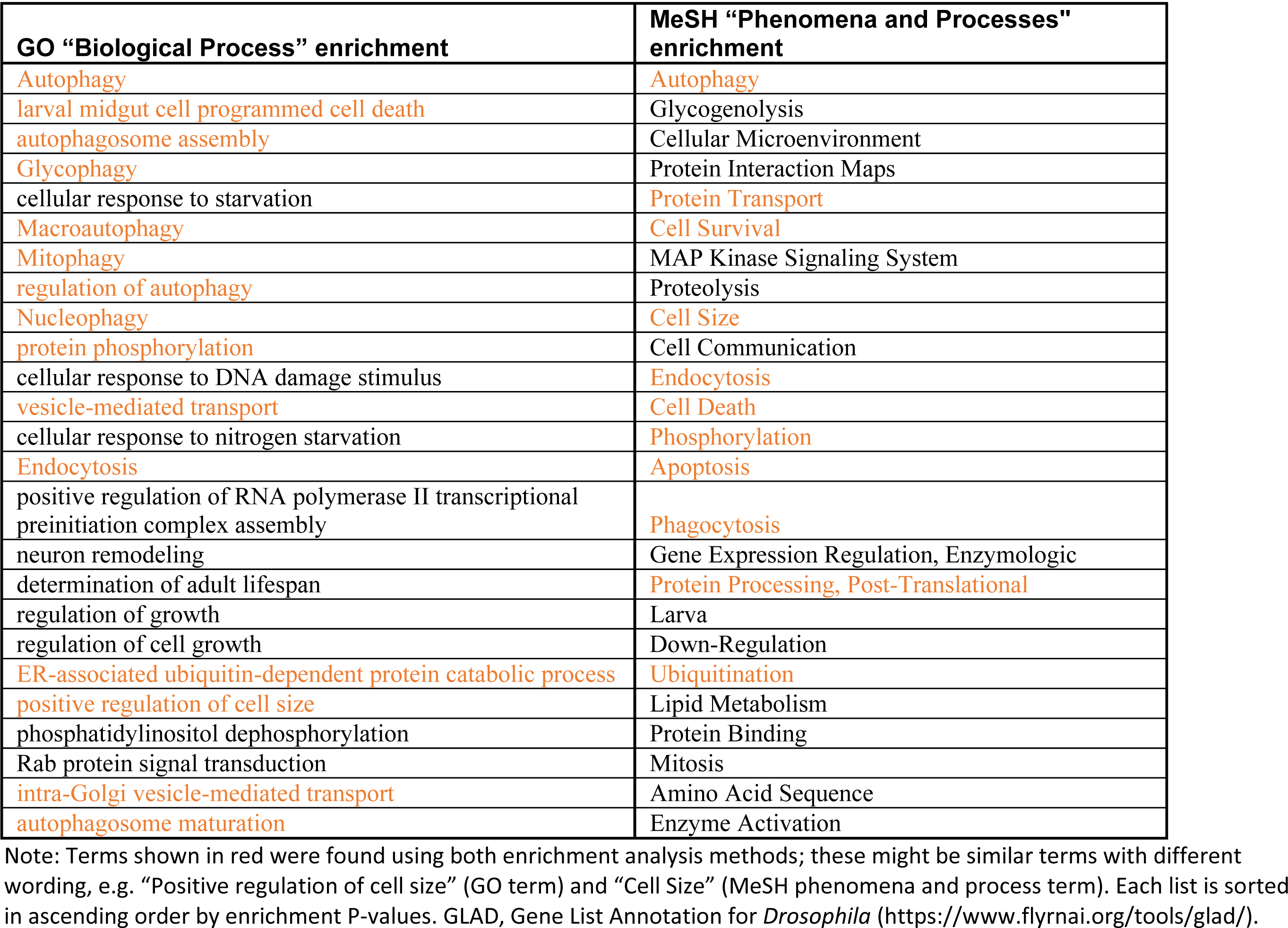
Comparison of the top 25 terms of GO enrichment analysis and MeSH enrichment analysis results for the set of autophagy genes provided at GLAD.

**Figure 5:**
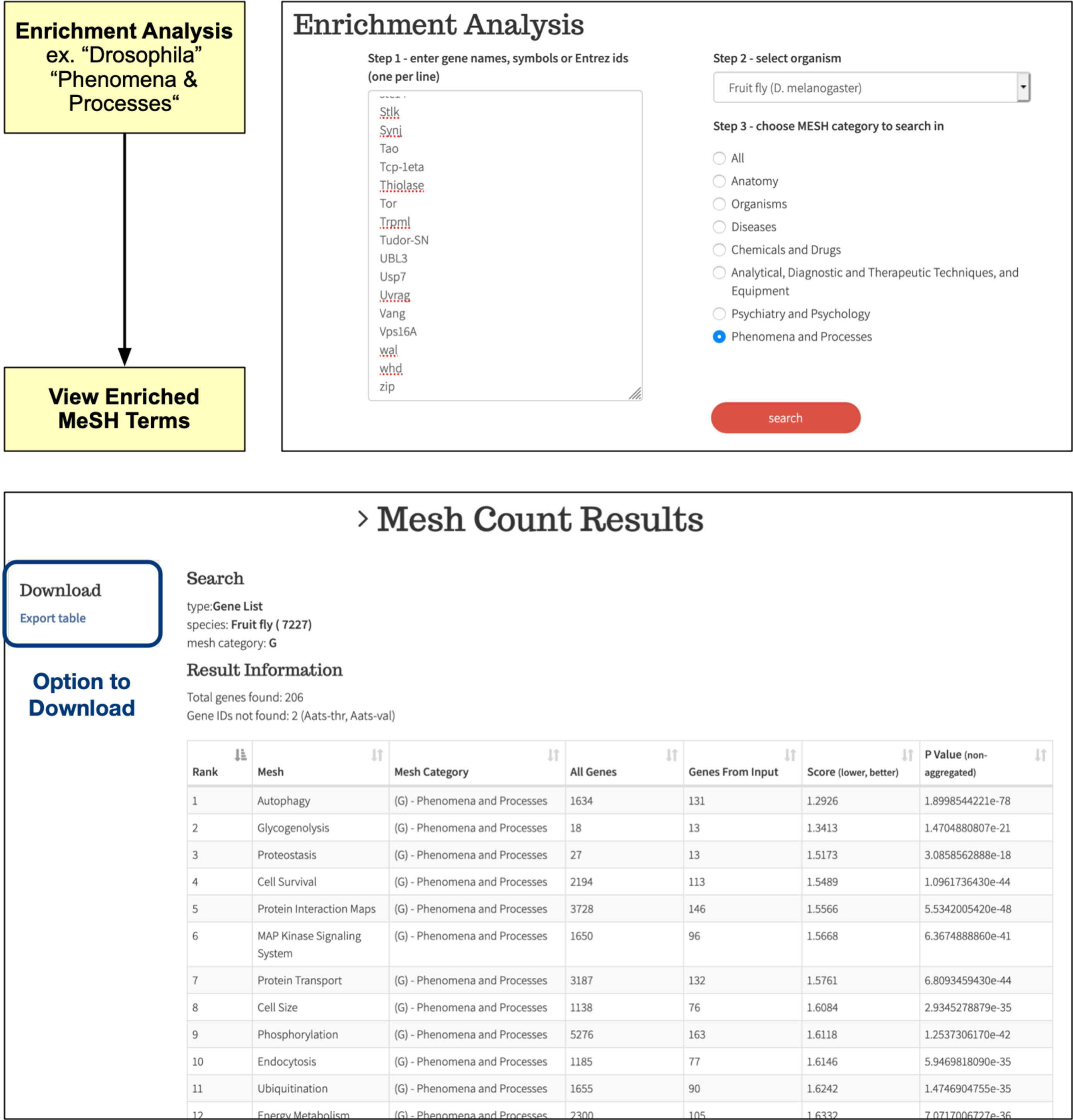
BioLitMine enrichment analysis. Example of an enrichment analysis of a gene list. In this example, the input is a list of 208 autophagy-associated genes from the Gene list Annotation for Drosophila (GLAD) online resource. In this example, the output as limited to MeSH terms in the category “phenomena and processes” (top panel). As expected, enrichment analysis identifies “autophagy” as a highly enriched term (bottom panel). The results can be downloaded as a csv file.

### Analysis of research trends

Using BioLitMine, we analyzed research trends represented by MeSH anatomy categories in publications for human, mouse, zebrafish and *Drosophila* studies (**Fig. 6**). The results of our analysis suggest that the MeSH anatomy terms “nervous system” and “urogenital system” are the most well-studied organs in *Drosophila*. This is in line with the long history of use of *Drosophila* to study neuroscience (” nervous system” term) and use of *Drosophila* ovaries and testes to study topics such as signal transduction and stem cell biology (” urogenital system” term). The anatomy terms “hemic/immune” and “nervous system” are the most represented in the gene-associated literature for human and mouse, whereas “embryonic structures” and “nervous system” are the most represented terms in the zebrafish literature.

**Figure 6.**
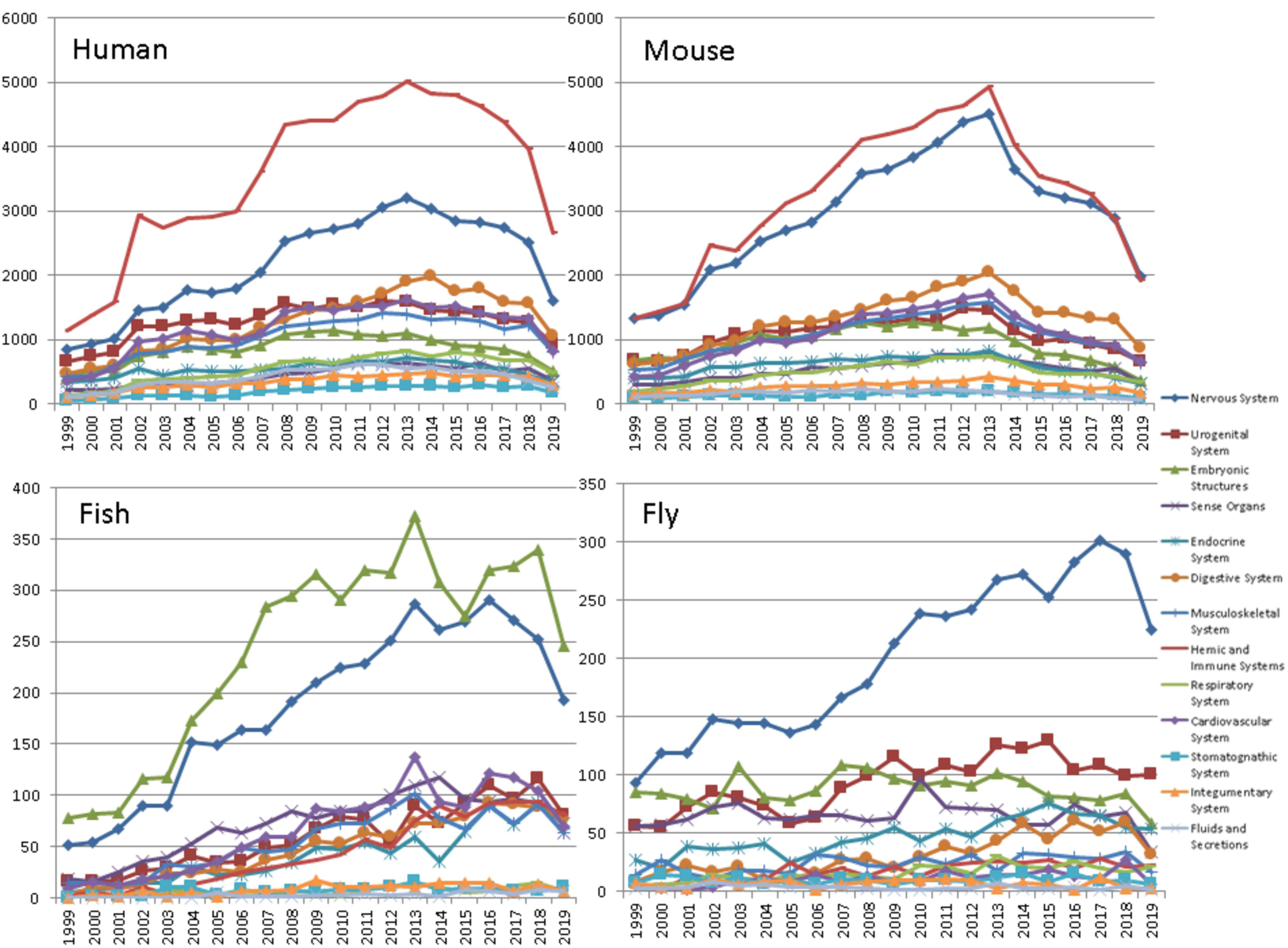
The trend of anatomy MeSH terms in MEDLINE publications for human, mouse, zebrafish and *Drosophila* studies. The count of gene-associated publications over time associated with anatomy MeSH terms for human, mouse, zebrafish and *Drosophila*. MeSH terms are organized in a hierarchical structure. For this analysis, we grouped publications based on root-level anatomy terms.

### Summary and conclusions

BioLitMine differs from similar resources in the following ways. 1.) BioLitMine provides a variety of search options for users; namely, gene-to-MeSH, MeSH-to-gene, gene-to-gene, gene-to-people, and pathway-to-people searches. In particular, the ability to do gene-to-people and pathway-to-people searches provide functions not well supported by other tools. This feature should make it easier for users to identify experts who might serve as scientific collaborators, reviewers, and so on, in a manner that is both automated and regularly updated. 2.) BioLitMine provides graphic visualization of search results. For example, it provides a word cloud for gene-to-MeSH search results, and a network view for gene-to-gene search results. 3.) BioLitMine compares co-citation networks with protein-protein interaction data, as well as genetic interaction data, using different edge colors to reflect the overlap. 4.) BioLitMine provides ortholog mapping options so that users can map genes identified from the literature associated with one species to conserved genes in another species, and retrieve relevant publications from both species. 5.) BioLitMine provides enrichment analysis of MeSH terms represented by all the related literature in an unbiased way, based on a user-inputted gene list.

Altogether, BioLitMine provides a powerful tool for literature mining that can help scientists gain a big picture of what the literature can offer for a gene of interest before diving into specific papers, and more specifically, can help scientists identify genes relevant to a MeSH term or another gene, in one or multiple species, as well as identify experts associated with a given pathway or gene.

## Methods

### Pipeline development

Our PubMed literature processing pipeline was developed to automatically retrieve pubmed2gene association files from NCBI and daily release files from the PubMed “pubmed ftp” site at the beginning of each month, as well as retrieve PubMed baseline files and MeSH annotation files at the beginning of each year. The pipeline then extracts the relevant information, such as information about authors and MeSH terms in the PubMed release files. It also precomputes all associations, such as gene-to-MeSH and gene-to-people associations, and updates the corresponding tables in the back-end mySQL database with this information. The pipeline was written in R, and Perl Scripts tied together with shell scripts and is hosted on the “O2” high-performance cluster made available by the Research Computing group (RC) at Harvard Medical School.

### Implementation of the web-based tool

The BioLitMine web tool (https://www.flyrnai.org/tools/biolitmine/) can be accessed directly or found at the ‘Tools Overview’ page at the DRSC/TRiP Functional Genomics Resources website (https://fgr.hms.harvard.edu/) (Hu *et al*. 2017). The back end was written using PHP with the Symfony framework. The front end HTML pages take advantage of the Twig template engine. The JQuery JavaScript library with the DataTables plugin is used for handling Ajax calls and displaying table views. The Bootstrap framework and some custom CSS are also used on the user interface. The website is hosted on the O2 high-performance computing cluster made available by the RC at Harvard Medical School. We use the third party software d3 (https://d3js.org/) and the Jason Daives d3-cloud package (https://github.com/jasondavies/d3-cloud) to create word clouds. The co-citation network was built with cytoscape.js (https://js.cytoscape.org/) (Franz *et al*. 2016).

### Data sources

NCBI PubMed base line files: https://ftp.ncbi.nlm.nih.gov/pubmed/baseline/

NCBI PubMed daily update files: https://ftp.ncbi.nlm.nih.gov/pubmed/updatefiles/

Gene2pubmed.gz file: https://ftp.ncbi.nlm.nih.gov/gene/DATA/

MeSH annotation files: https://www.nlm.nih.gov/databases/download/mesh.html

## Acknowledgements

We would like to thank the members of Perrimon laboratory, FlyBase consortium, Drosophila RNAi Screening Center (DRSC) and Transgenic RNAi Project (TRiP) for helpful input on the project. Relevant grant support includes NIH NIGMS P41 GM132087. In addition, S.E.M. is supported in part by the Dana Farber/Harvard Cancer Center, which is supported in part by NIH NCI Cancer Center Support Grant P30 CA006516. N.P. is an investigator of Howard Hughes Medical Institute.

## Availability

https://www.flyrnai.org/tools/biolitmine/web/

